# Increased protein expression of methylenetetrahydrofolate reductase and cystathionine β-synthase in medial prefrontal cortical tissue of female vascular dementia patients

**DOI:** 10.1101/2025.11.21.689865

**Authors:** Sanika M. Joshi, Abbey McKee, Sharadyn Ille, Kristina Buss, Thomas G Beach, Geidy E Serrano, Nafisa M. Jadavji

## Abstract

Vascular dementia (VaD) is a complex clinical syndrome arising from cerebrovascular disease, characterized by cognitive decline and functional impairment, and is projected to double in prevalence over the next three decades. Deficiencies in one-carbon (1C) metabolism are linked to the onset of VaD. Our previous work using mouse models has demonstrated that reduced dietary folic acid intake or genetic disruptions in 1C metabolism exacerbate outcomes in a model of VaD. However, the impact of VaD on one-carbon metabolism remains poorly understood. This study aims to provide a detailed molecular portrait of 1C metabolism within the context of VaD, shedding light on potential molecular mechanisms. In post-mortem medial prefrontal cortex tissue from VaD female and male patients and controls we measured protein expression of the folate receptor (FR) and 1C enzymes including methylenetetrahydrofolate reductase (MTHFR), thymidylate synthase (TS), choline acetyltransferase (ChAT), acetylcholinesterase (AChE), cystathionine β-synthase (CBS) co-localized with NeuN. There was an interaction between VaD and gender for levels of FR. Both male and female VaD had increased levels of ChAT. Female VaD patients had higher levels of MTHFR and CBS when compared to males. VaD is a complex disease; the results of this study demonstrate that VaD impacts neuronal levels of 1C enzymes. Future studies should assess 1C in other cell types of the brain, as well as measure enzyme activity levels.

**Graphical Abstract:** 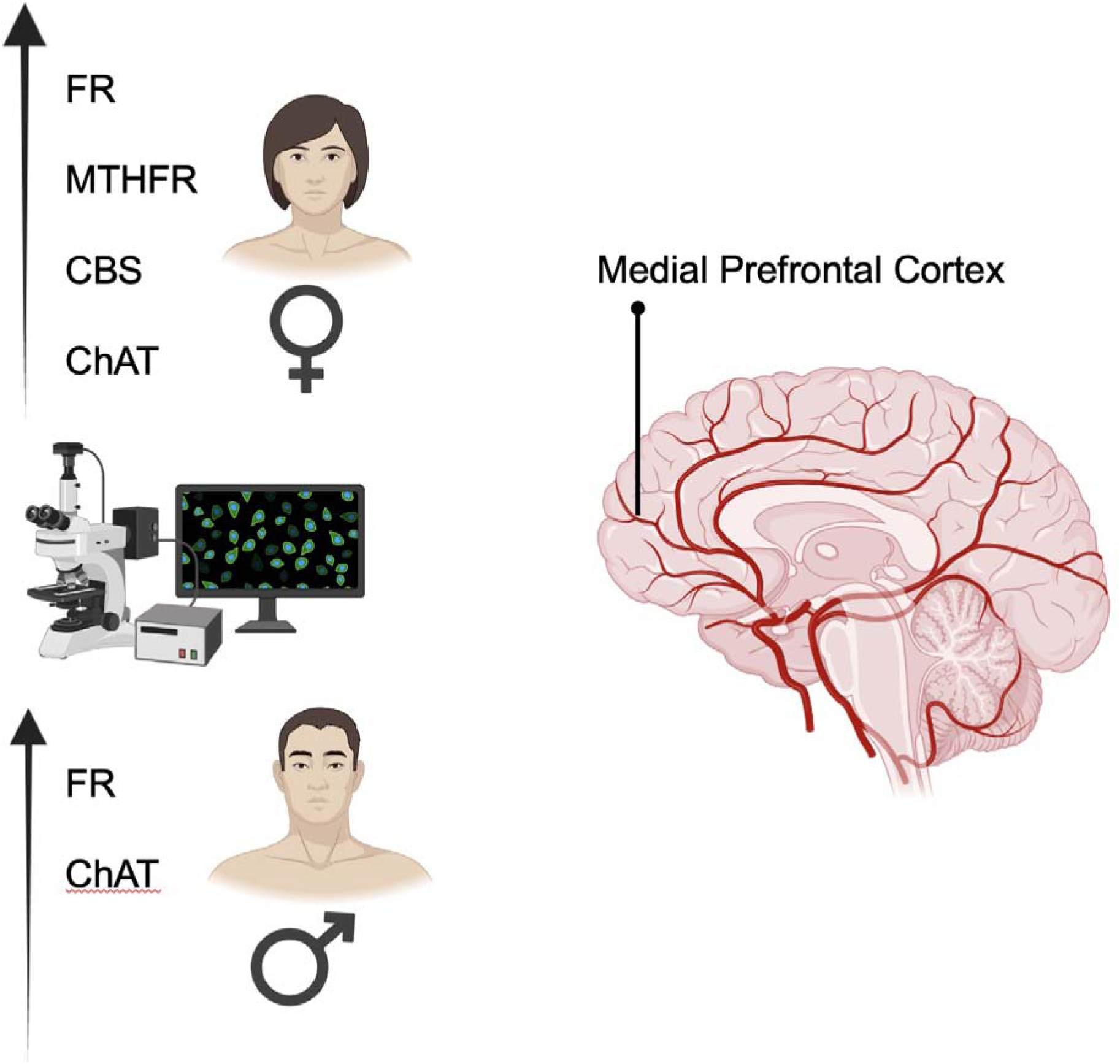

## Introduction

Vascular dementia (VaD) is projected to double in prevalence within the next three decades and represents a complex and multifaceted clinical syndrome arising from cerebrovascular disease, characterized by cognitive decline and functional impairment (Gorelick et al., 2011). VaD comprises a spectrum of cognitive disorders resulting from cumulative vascular pathologies affecting the brain. The cumulative burden of cerebrovascular lesions, especially intracranial small vessel disease (SVD), correlates with VaD, featuring common occurrences of lacunar infarcts, microinfarcts, and diffuse white matter changes (Kalaria, 2016).

Several subtypes of VaD have been identified, highlighting the complexity of the relationship between vascular issues and cognitive decline (O’Brien et al., 2003). One notable aspect of VaD is that there is no single neuropsychological pattern that can definitively distinguish it from other causes of cognitive impairment. The underlying vascular pathology of VaD shares pathogenic elements, including hypoperfusion, increased oxidative stress and inflammation, leading to endothelial damage, blood-brain barrier breakdown, innate immunity activation, and disruption of trophic coupling between vascular and brain cells (Iadecola, 2013). Nutritional status has been linked to VaD (Smith et al., 2018).

Deficiencies in one-carbon (1C) metabolism, such as dietary deficiencies in folic acid or choline have been linked to cognitive impairment, including VaD in the elderly population (Spence, 2016; Smith et al., 2018). Our work has shown that there is a complex interplay between dietary, genetic, and vascular factors in the manifestation of cognitive deficits (Jadavji et al., 2015; Dam et al., 2017; Joshi and Jadavji, 2024). We have demonstrated novel insights into arterial remodeling, GFAP expression, and MMP-9 levels, contributing to a more nuanced understanding of the pathophysiological mechanisms underlying VaD (Jadavji et al., 2015). These findings underscore the significance of considering the multifaceted nature of VaD. Another study from our group challenged existing controversies surrounding the association between folic acid metabolism, homocysteine, and VaD, shedding light on the nuanced impact of these factors on cognitive function, vasculature, and choline metabolism (Dam et al., 2017). Understanding of the role of 1C metabolism in VaD requires further investigation. The aim of this study was to investigate the levels of 1C enzymes and receptors in post-mortem brain tissue from female and male patients diagnosed with VaD.

## Methods

### Brain tissue samples

This study was conducted in accordance with the Declaration of Helsinki and was determined to be Internal Review Board (IRB) exempt as per the Midwestern University IRB Committee. Human cortical brain tissue was obtained from Banner Sun Health Research Institute Brain and Body Donation Program (Beach et al., 2015). Informed consent was obtained from all individual participants included in the study. Samples were obtained from postmortem female and male patients with a vascular dementia diagnosis and healthy controls. Details of samples including age and comorbidities are listed in **Table 1**.

**Table 1.** Patient sample demographics and comorbidities.

|  | **Control** |  | **VaD** |  |
| --- | --- | --- | --- | --- |
| **Age** | **Male** | **Female** | **Male** | **Female** |
| >90+ | 1/4 (25%) | 3/4 (75%) | 0/4 (0%) | 3/4 (75%) |
| 80 to 90 | 3/4 (75%) | 1/4 (25%) | 4/4 (100%) | 1/4 (25%) |
| Vascular Dementia | 0/4 | 0/4 | 4/4 | 4/4 |
| Cerebral white matter rarefaction | 0/4 | 0/4 | 4/4 | 4/4 |
| **Comorbidities** |  |  |  |  |
| CAA | 2/4 (50%) | 2/4 (50%) | 0/4 (0%) | 1/4 (13%) |
| HR | - | - | 0/4 (0%) | 1/4 (13%) |
| LBS | - | - | 2/4 (50%) | 1/4 (13%) |

Abbreviations: CAA - Cerebral amyloid angiopathy; HR - Hippocampal Sclerosis, LBS- Lewy Bodies; VaD – vascular dementia

### Immunofluorescence Experiments

To investigate 1C enzyme levels co-expression within neuronal nuclei, immunofluorescence experiments of VaD and control brain tissue was performed. Several primary antibodies were used to identify the specific protein in question for a given sample: FR (1:100, Thermo Fisher, Waltham, MA, USA, RRID: AB_2609390), MTHFR (1:100, AbCam, Boston, MA, USA, RRID: AB_2893493), ChAT (1:100, Millipore Sigma, Darmstadt, Germany, RRID: AB_2079751), AChE (1:100, Millipore Sigma, Darmstadt, Germany, RRID: AB_10602654), CBS (1:100, ThermoFisher, catalog number: MA5-17273) were used. All brain sections were stained with a marker for neuronal nuclei, NeuN (1:200, AbCam, Waltham, MA, USA, RRID: AB_10711040). Primary antibodies were diluted in 0.5% Triton X and incubated on the brain tissue overnight at 4°C. Brain sections were incubated with secondary antibodies Alexa Fluor 488 or 555 (1:200, Cell Signaling Technologies) the following day at room temperature for 2h.

For each antibody we used an n = 3 samples per group, as determined by *a priori* sample size calculations using G*3 Power (Faul et al., 2007). The observed effect size was d=1.25, alpha level was 0.05, power was set to 80% (Bacchetti, 2010). If there was variability staining, the sample size was increased to 4.

### Immunofluorescence Analysis

Microscope slides were cover slipped with fluromount and stored at 4 until analysis. The staining was visualized using the Echo Revolve microscope (Echo, San Diego, CA, USA) and all images were collected at the magnification of 200×. To assess 1C protein levels within ischemic stroke and control brain tissue, colocalization of the primary antibody with NeuN and DAPI-labeled neurons was counted and averaged across three sections per primary antibody per subject. Cells were distinguished from debris by identifying a clear cell shape and intact nuclei (indicated by DAPI and NeuN) under the microscope. All cell counts were conducted by two individuals blinded to treatment groups. Using ImageJ, the number of positive cells was counted for a total of 8 healthy aged control subjects (4 males, 4 females) and 8 VaD subjects (4 males, 4 females).

### Statistics

GraphPad Prism 6.0 was used to immunofluorescence staining quantification. In GraphPad Prism 10.6.0, D’Agostino-Pearson normality test was performed prior to two-way ANOVA analysis when comparing the mean measurement of both sex and group (control or VaD). Significant main effects of two-way ANOVAs were followed up with Tukey’s post-hoc test to adjust for multiple comparisons. All data are presented as mean ± standard error of the mean (SEM). Statistical tests were performed using a significance level of 0.05.

## Results

### Folate Receptor

Using immunofluorescence, we stained brain tissue from healthy controls and VaD patients for the folate receptor (FR). Representative staining from all groups of the FR are shown in **Figure 1A**. There was a significant interaction between VaD and gender (**Figure 1B**; F(_1,11_) = 5.41, p = 0.040). There was no difference between genders (p = 0.27) or disease state (p = 0.49).

**Figure 1.**
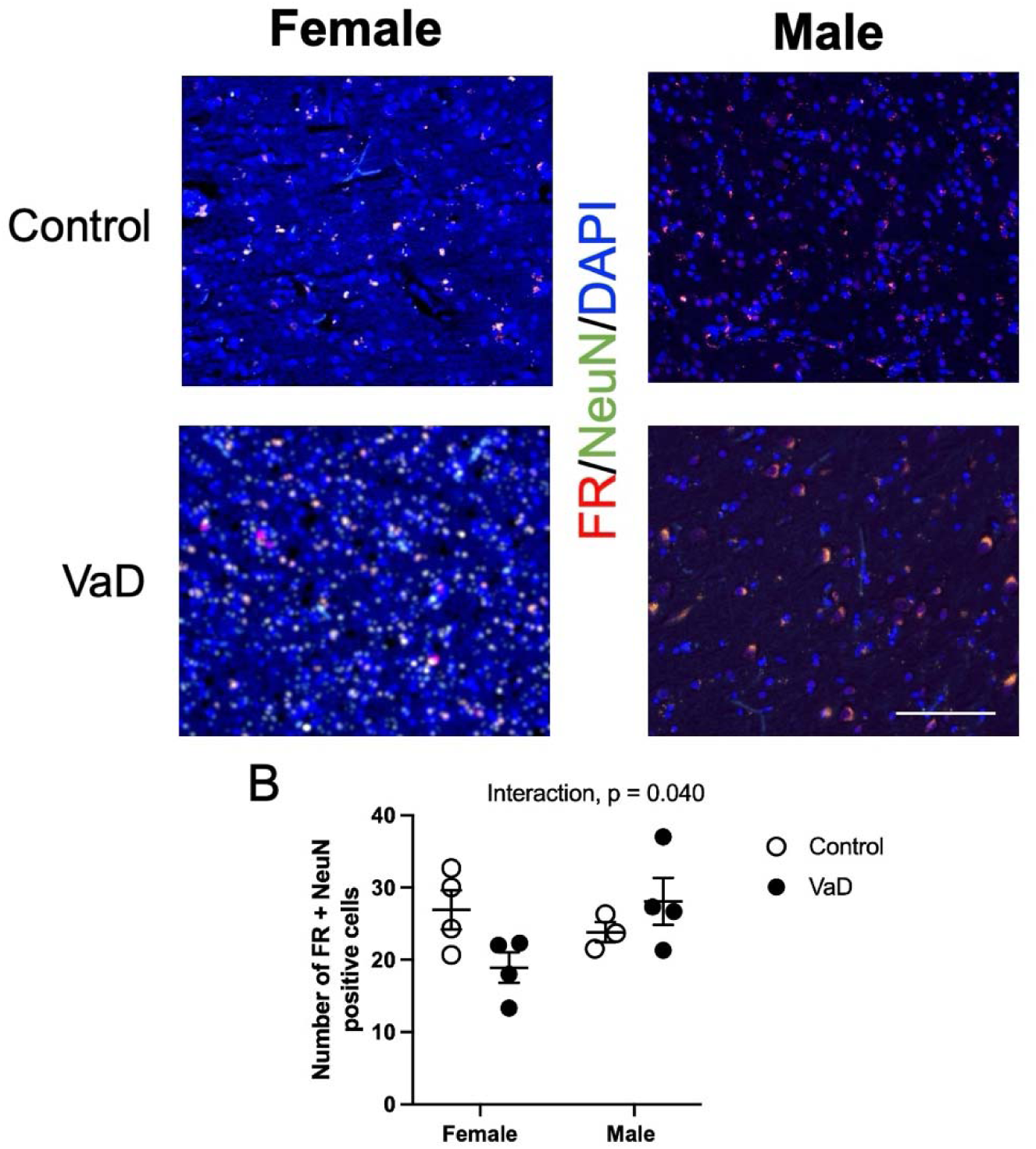
Levels of folate receptor (FR), co-stained with neuronal nuclei (NeuN) and DAPI in brain tissue of controls and Vascular Dementia (VaD) patients. Representative images (A) and quantification of levels (B). Significant interaction between disease and sex, p < 0.05. Data represents 4 patients per group. Scale bar at 50 μm.

### Methylenetetrahydrofolate reductase (MTHFR)

Representative staining from all groups of methylenetetrahydrofolate reductase (MTHFR) are shown in **Figure 2A**. MTHFR expression in cortical tissue was impacted by VaD (**Figure 2B**, F(_1,12_) = 5.41, p = 0.023). Female VaD patients had higher levels of MTHFR in cortical tissue compared to healthy controls (p = 0.0092). There was no interaction (p = 0.09) or impact of gender (p = 0.60) on MTHFR levels. We checked for outlier using the GraphPad Quick Cals Grubb’s test. The z was 1.42267 when the p-value was set as 0.05, therefore there was no significant outlier.

**Figure 2.**
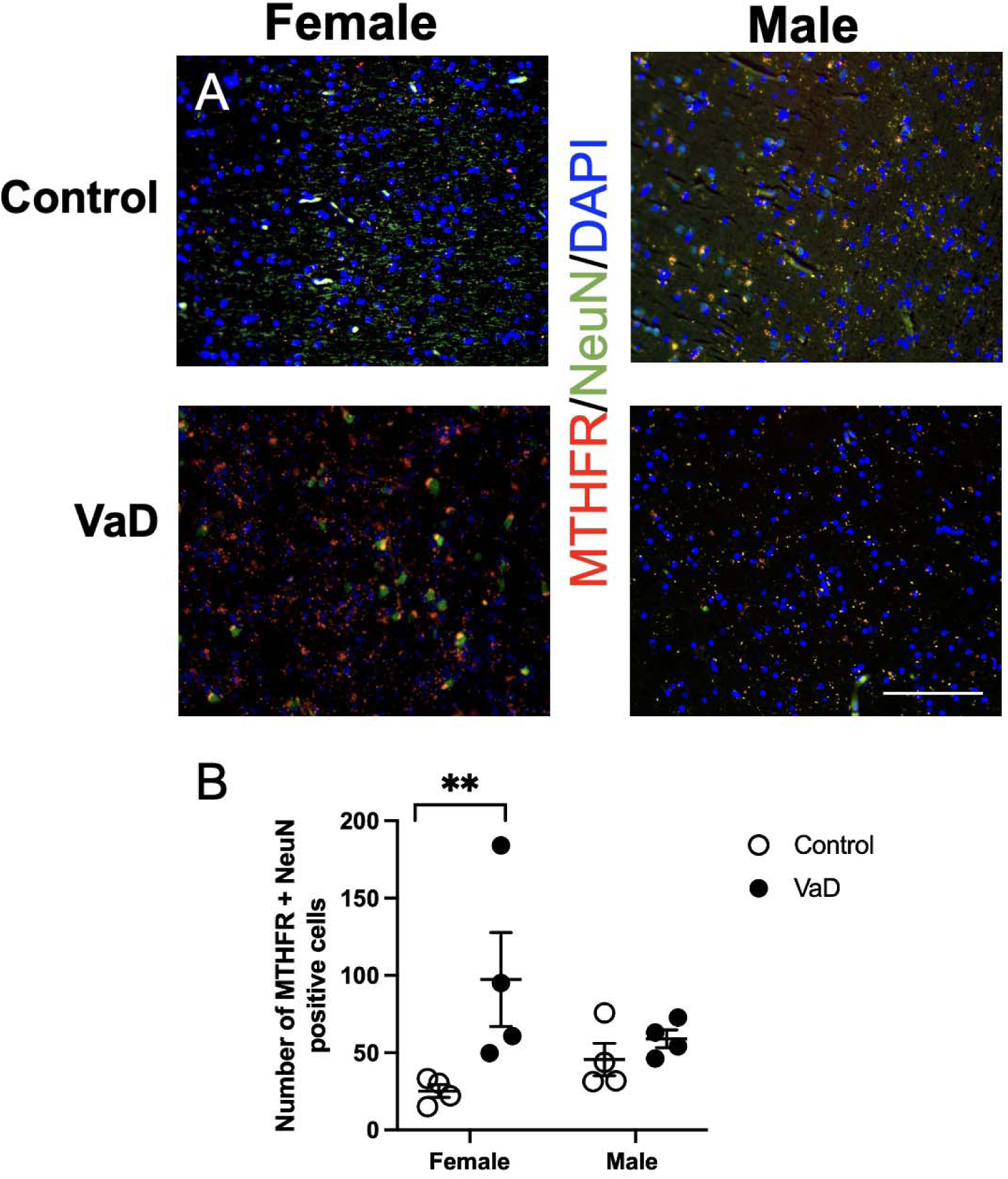
Levels of methylenetetrahydrofolate reductase (MTHFR), co-stained with neuronal nuclei (NeuN) and DAPI in brain tissue of controls and Vascular Dementia (VaD) patients. Representative images (A) and quantification of levels (B). Data represents 4 patients per gender, per group. ** p < 0.01, Tukey’s pairwise comparison between female control and VaD patients. Scale bar at 50 μm.

### Choline acetyltransferase (ChAT)

Representative staining from all groups of choline acetyltransferase (ChAT) are shown in **Figure 3A**. ChAT protein expression were increased in VaD patients (**Figure 3B**, F(_1,12_) = 4.95, p = 0.046). There were no differences between groups. There was no interaction (p = 0.75) or impact of gender (p = 0.46) on ChAT levels.

**Figure 3.**
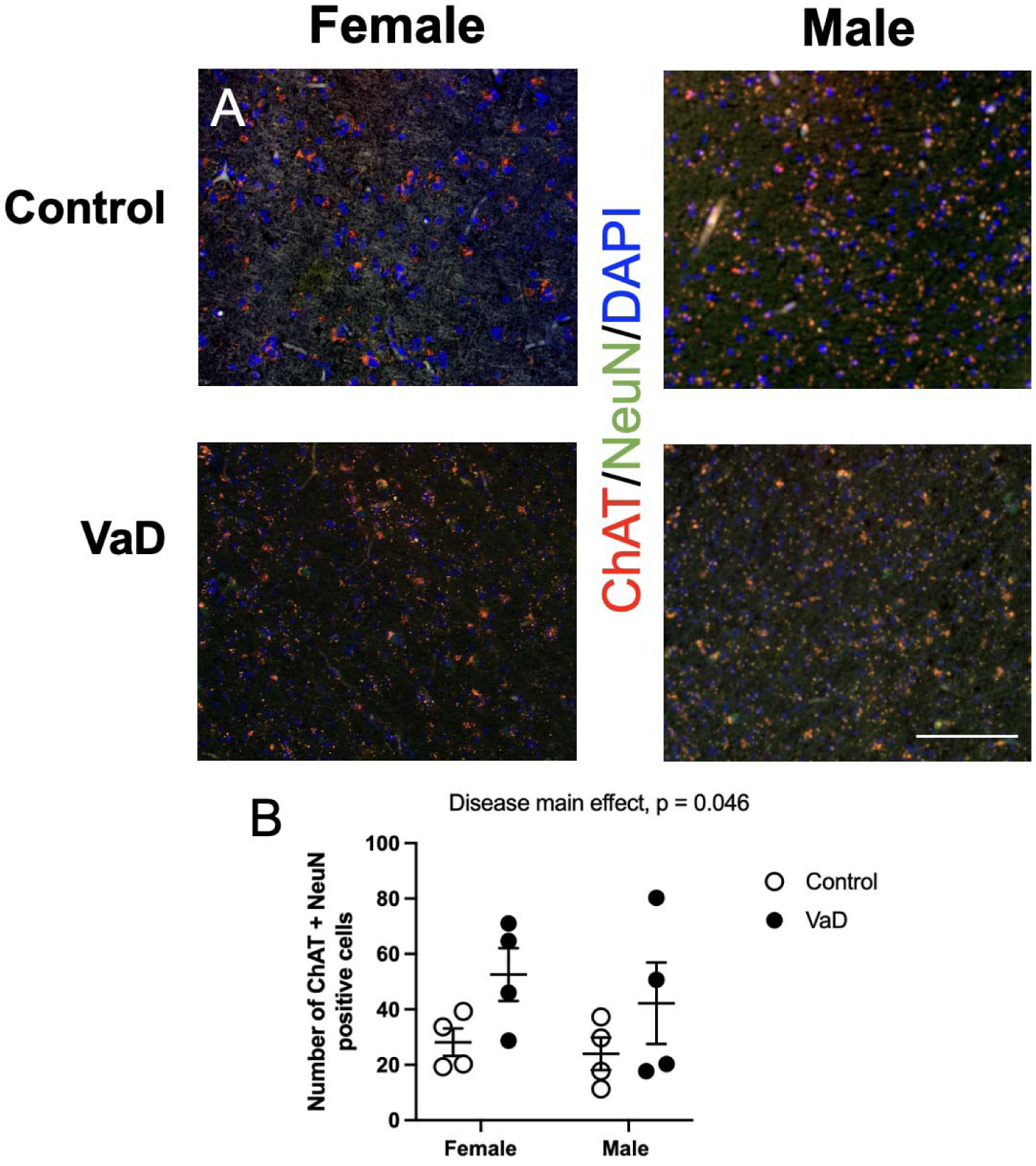
Levels of choline acetyltransferase (ChAT), co-stained with neuronal nuclei (NeuN) and DAPI in brain tissue of controls and Vascular Dementia (VaD) patients. Representative images (A) and quantification of levels (B). Data represents 4 patients per group. Scale bar at 50 μm.

### Acetylcholinesterase (AChE)

AChE levels were not impacted by VaD (p = 0.45, data not shown) or gender (p = 0.41,). There was no interaction between VaD and gender (p = 0.44).

### Cystathionine-***β***-synthase (CBS)

Representative staining of cystathionine-β-synthase (CBS) is shown in **Figure 4A**. CBS protein expression was impacted by gender (**Figure 4B**, F(_1,11_) = 7.32, p = 0.0204). VaD female had higher levels compared to male patients (p = 0.033). There was no interaction (p = 0.41) or impact of VaD (p = 0.0585) on CBS expression.

**Figure 4.**
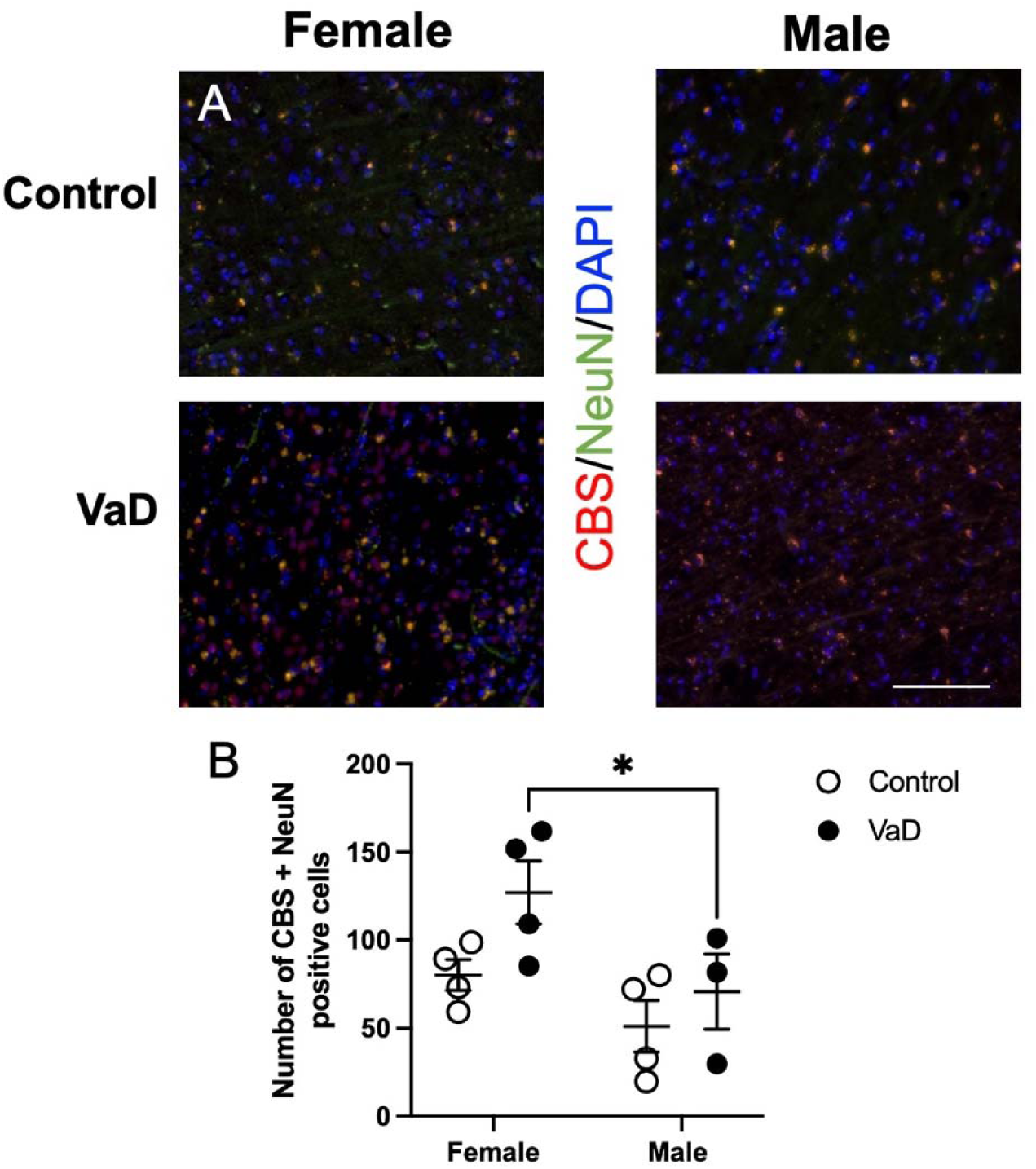
Levels of Cystathionine-ß-synthase (CBS) co-stained with neuronal nuclei (NeuN) and DAPI in brain tissue of controls and Vascular Dementia (VaD) patients. Representative images (A) and quantification of levels (B). Data represents 4 patients per group. * p < 0.05, Tukey’s pairwise comparison between female control and VaD patients. Scale bar at 50 μm.

## Discussion

VaD comprises a spectrum of cognitive disorders resulting from cumulative vascular pathologies affecting the brain. It can be challenging to identify the clinical and pathological substrates of cerebrovascular disease in the context of VaD, with vascular changes often found in the elderly either alone or alongside neurodegenerative processes (Kalaria, 2016). Additionally, several subtypes, highlighting the complexity of the relationship between vascular issues and cognitive decline (O’Brien et al., 2003). To further elucidate the intricate relationship between one-carbon (1C) metabolism and vascular dementia (VaD), the present study aimed to investigate the protein levels of 1C enzymes and the folate receptor in post-mortem medial prefrontal cortical brain tissue from VaD and healthy aged patients. Our analysis shows that the folate receptor levels were impacted by VaD and gender. Female VaD patients had increased levels of MTHFR and CBS. VaD impacted choline metabolism only through increased levels of ChAT in female and male patients. ChAT makes acetylcholine at synapses. Reports show that there are reduced levels of acetylcholine in VaD patients, which corresponds to increased levels of ChAT in cortical brain tissue (Parnetti et al., 2007). Our previous work (Jadavji et al., 2012; Dam et al., 2017), as well as others (Wurtman, 1992) has demonstrated reduced levels of acetylcholine result in increased levels of ChAT. Our work has provided some insights into how 1C is impacted in the VaD disease state.

Choline is an essential nutrient found in food that is a precursor in the synthesis of phospholipids, betaine, and acetylcholine. Betaine is a methyl group donor for homocysteine to methionine reaction. Acetylcholine is an important neurotransmitter involved in neurogenesis, synapse formation, learning and memory (Blusztajn, 1998). Changes in choline metabolism have been reported in VaD (Erkinjuntti et al., 2004; Wang et al., 2009; Blusztajn et al., 2017). Our work confirms these changes through immunofluorescence experiments. Choline is made endogenously in our body, but we require dietary supplementation – and the neuroprotective effects of increasing choline have been shown by our group using a mouse model of ischemic stroke (Jadavji et al., 2017, 2019), as well as others (Blusztajn et al., 2017). Recent reports show that ∼90% of the US population does not reach the acceptable daily intake (ADI) of choline (Zeisel, 2017). Recent work has demonstrated that adequate dietary choline intake is necessary to reduce dementia hallmarks (Dave et al., 2023).

To follow up clinical findings that increased levels of homocysteine were linked to cognitive impairment (Seshadri et al., 2002), several research groups have shown that changes in 1C impact cognitive impairment (Hainsworth et al., 2015; Jadavji et al., 2015; Gooch and Wilcock, 2016; Dam et al., 2017; Joshi and Jadavji, 2024). In our opinion investigating the link of 1C and VaD in post-mortem brain tissue was the next step. Our present study shows that the FR and MTHFR, ChAT, and CBS enzymes are impacted by VaD. Interestingly there is also a sex difference, which have also been reported in the clinical population (Akhter et al., 2021). We measured levels of enzymes in prefrontal medial cortical tissue, investigating other areas of the brain as well as other cell types in the brain, glial and endothelial cells would provide more details. Investigation of 1C markers in blood, such as homocysteine would also provide more data.

Homocysteine is marker of 1C metabolism as well as disease state (Joshi and Jadavji, 2024). Clinical studies have revealed a relative risk of dementia in the elderly with moderately raised homocysteine within the normal range, ranging from 1.15 to 2.5, and a Population Attributable risk from 4.3% to 31% (Smith et al., 2018). Intervention trials with B vitamins in elderly individuals with cognitive impairment demonstrate a significant slowing of whole and regional brain atrophy, along with a deceleration of cognitive decline (Smith et al., 2010). These findings suggest that moderately raised plasma total homocysteine (>11 mol/L), prevalent in the elderly, may contribute to age-related cognitive decline and dementia. The International Consensus Statement, drawn from a comprehensive review of literature spanning the last two decades, emphasizes the importance of identifying modifiable risk factors for the prevention of dementia (Smith et al., 2018). The public health significance of addressing elevated homocysteine is underscored, given its ease, cost-effectiveness, and safety of treatment with B vitamins. In 2024, the Food for the Brain foundation launched a home test to measure homocysteine levels in the elderly, with the goal to catch impaired 1C metabolism and correct it with dietary supplementation (Anon, n.d.). Further trials are recommended to explore whether B vitamin treatment can slow or prevent the conversion to dementia in individuals at risk of cognitive decline or dementia.

## Conclusions

VaD is a complex disease; the results of this study demonstrate the impact of VaD on 1C and changes in gene expression. Supplementation with 1C in VaD patients may be beneficial for VaD affected patients as well as targeting reduced gene expression to increase better outcomes.

## Credit authorship contribution statement

Conceptualization, N.M.J.; methodology, N.M.J., S.M.J., A.M, K.B., T.G.B., G.E.S, ; /software, X.X.; validation, N.M.J.; formal analysis, S.M.J., A.M, S.I, K.B., N.M.J.; investigation, S.M.J., A.M, S.I, K.B., N.M.J.; resources, N.M.J.; data curation, S.M.J., A.M, S.I, K.B., N.M.J.; writing—original draft preparation, S.M.J., K.B., N.M.J.; writing—review and editing, X.X.; visualization, N.M.J.; supervision, N.M.J.; project administration, N.M.J., T.G.B., G.E.S.,; funding acquisition, N.M.J., T.G.B., G.E.S. All authors have read and agreed to the published version of the manuscript.

## Declaration of generative AI use

The authors report generative AI was not used in their research or preparation of this manuscript.

## Data Availability Statement

Data is available on Dryad, DOI: 10.5061/dryad.r7sqv9srv

